# The Role of Rivers as Geographical Barriers in Shaping Genetic Differentiation and Diversity of Neotropical Primates

**DOI:** 10.1101/2023.07.23.550208

**Authors:** William D. Helenbrook, Jose Valdez

## Abstract

We quantitatively tested the riverine barrier hypothesis and its influence on biogeographical distributions and molecular variation in New World monkeys (Parvorder: Platyrrhini). Using mitochondrial markers (cytochrome oxidase subunit II and cytochrome b), we analyzed taxonomic differences and the effects of geographical barriers on molecular patterns across Central and South America. Nearly 80% of described species are separated by geographical barriers. River width exhibited a positive correlation with molecular dissimilarity in adjacent taxa for both molecular markers. Streamflow also showed a positive association, although not statistically significant, likely due to limited sample size. Several presently described taxa were not supported based solely on these molecular phylogenetic markers, including *Saimiri*, *Mico*, *Cebus*, *Sapajus*, and *Cherecebus*. These taxonomic issues are far more common where river barriers do not exist. In conclusion, we found a significant correlation between river width and molecular divergence in adjacent taxa, indicating that wider rivers were associated with greater molecular dissimilarity for two commonly used mitochondrial genes. Species boundaries were predominantly found at river interfaces, and in the absence of discernable geological barriers, adjoining species were more likely to exhibit molecular similarity. Our findings suggest that both river and mountain barriers significantly restrict gene flow for the majority of neotropical taxa, with geological formation of river barriers coinciding with estimated speciation events. Additionally, river width proves to be a valuable tool for estimating molecular divergence in adjacent taxa, particularly in regions with limited sampling.

## 1. INTRODUCTION

The Neotropics, a region encompassing Central America, South America, and the Caribbean islands, contains approximately one-third of all known primate species (Paglia 2012). This region is home to a high concentration of unique and native species and is predicted to be an area of high potential for future discoveries (Moura & Jetz, 2021). This is due to its large diversity of habitats attributed to its geological and climatic history, resulting in complex ecological barriers driven by diverse habitats and elevational gradients. The retracted and fragmented forests in the Amazon basin, resulting from glacial advances during Pleistocene climatic fluctuations and Pliocene-Miocene orogenic events, has presumably played a critical role in the evolution of these distinct areas of endemism in accordance with the Refugia hypothesis (Haffner 1969; Prance, 1982; Turchetto-Zolet et al. 2013; Thom et al. 2020), though the degree to which this mechanism has driven Amazon diversity on its own has been challenged (Bush and Oliveira 2006; Rocha and Kaefer, 2019). The question remains whether climate fluctuations contributed to either retraction and expansion of stable forested habitats, or whether areas of endemism were isolated by climate driven fluvial barriers across the Amazon basin, ultimately contributing to the considerable diversity of New World monkeys (parvorder: Platyrrhini) (Chapman, 1926; Prance, 1982; Burney and Brumfield, 2009).

The river barrier hypothesis, proposed by Alfred Wallace in the 19th century hypothesizes that rivers limit or even prevent the dispersal of populations, leading to isolation, and ultimately to divergence through the differential accumulation of mutations, local adaptation, and genetic drift; a process of speciation known as vicariance (Wallace, 1852; Wallace, 1876; Lomolino et al., 2010). The river barrier hypothesis proposes that landscape-level drainage systems can limit the distribution of taxa (Ayers and Clutton-Brock 1992) and supported by recent geochronological approaches (Pupim et al. 2019; Mourthe et al. 2022), suggesting that shifting semi-permeable barriers might enable sporadic gene flow, while strong river barrier formation restricts gene flow and subsequent speciation (Ribas et al. 2012; Musher et al. 2022). The complexity of successional fluvial deposition and fluctuating landscape connectivity across geological time explains why some biogeographical patterns are distinctly associated with riverine barriers, while in other cases the relationship between geological history and taxonomic diversification is more complicated. Understanding the different factors that contribute to the high diversity of species in the neotropics and how their influence varies across taxonomic groups is crucial.

In the Amazon basin, the distribution of New World monkey species has been found to be strongly correlated with the distribution of rivers. Some species are found only on one side of a river or other body of water, providing evidence for the role of these waterbodies driving speciation (Lomolino et al., 2010). This is also supported by genetic studies which have revealed high levels of genetic variation within populations and low levels of variation between populations separated by rivers, consistent with the idea that these species have evolved independently (Beaudrot & Marshall, 2011; Harcourt & Wood, 2012). Evidence of rivers restricting gene flow and leading to differentiation in neotropical primates has also been reported in several genera including *Saimiri* (Chiou et al., 2011; Alfaro et al., 2015; Ruiz-Garcia et al., 2015), *Alouatta* (Cortés-Ortiz et al., 2003; Ruiz-Garcia et al., 2017), *Sapajus* (Martins-Junior et al., 2018; Boubli et al. 2019), *Lagothrix* (Ruiz-Garcia et al., 2019), *Cacajao*, *Callicebus*, *Cebus*, *Pithecia*, and *Saguinus* (Boubli et al., 2015).

The restriction of gene flow due to rivers is influenced by three key factors: the geographical history of river systems, dispersal ability of organisms, and river characteristics like size and speed. Firstly, molecular divergence is dependent upon the origin of the river and how long the barrier has restricted gene flow. For example, recent dynamic tributary rearrangements have been shown to influence biogeographical observations (Ruokolainen et al. 2019). The dispersal ability of the organism also plays a role, with larger, more mobile species being more likely to cross rivers (Lomolino et al., 2010). Furthermore, the restriction of dispersal caused by river barriers such as width, volume, and flow rate are positively also associated with divergence. This has been observed in primate species living on opposing sides of Amazonian rivers, where river strength and width are inversely correlated with species similarity (Ayres & Clutton-Brock, 1992; Fordham et al., 2020). However, despite the evidence in support of the river barrier hypothesis, further research is needed to fully understand how rivers and other bodies of water affect gene flow and speciation, and how other factors may also play a role in the process.

One approach to studying the impact of riverine barriers on the speciation of New World primates is to analyze partial mitochondrial genes, cytochrome b (cytb), and cytochrome oxidase subunit II (COII) genomes. These genes are widely used to trace evolutionary relationships and genetic diversity among populations and species. Mitochondrial DNA can provide insights into evolutionary relationships and the timing of divergence. We can compare molecular similarity – an indicator of gene flow between adjacent species on opposing riverbanks – with measures of river permeability in order to model the impact of these barriers on taxonomic divergence. Furthermore, quantifying the correlation between river width and flow rate with molecular similarity could potentially predict divergence in under-sampled areas. This information can aid in identifying areas of taxonomic significance, which can then be further explored through molecular, behavioral, and morphological analyses.

Mitochondrial DNA analysis serves as a powerful tool for studying elusive species like New World primates, where obtaining high-quality tissue samples through non-invasive methods is challenging. However, the full potential of mitochondrial markers is often hindered by limitations, including incomplete databases and the absence of crucial sampling details such as precise locations, posing obstacles in population comparisons. A recent study using mitogenomic data revealed that while some divergences coincided with river barriers, not all of them occurred simultaneously or aligned with proposed river formation dates (Janiak et al. 2022). The strongest evidence of restricted dispersal was found on opposing sides of the Amazon River, particularly in bearded saki monkeys (*Chiropotes* spp.) and marmosets/tamarins (small platyrrhines), with limited evidence for the Rio Negro. Surprisingly, large rivers did not act as barriers for certain primate species, including howler monkeys (Alouatta spp.), uakaris (Cacajao spp.), sakis (Pithecia spp.), and robust capuchins (Sapajus spp.). These findings suggest the involvement of other evolutionary mechanisms in the diversification of platyrrhine primates. While the study’s findings provide valuable insights, its limitations, such as a small sample size, lack of quantitative analysis, and focus solely within the Amazon basin, highlight the importance of conducting further research to deepen our understanding of how riverine barriers contribute to the diversification of platyrrhine primates.

In this study, we conduct a quantitative molecular analysis of the riverine barrier hypothesis on all new world monkeys across Central and South America. The main goal of the study is to investigate the role of geographical barriers, particularly rivers, in shaping the distribution and divergence of Platyrrhine primates in South America. The specific aims are to: 1) identify geographical boundaries such as river and mountain barriers while analyzing molecular phylogenetics of Platyrrhini across identified barriers 2) assess the relationship of river width and flow rate to molecular intra and interspecies diversity on opposing riverbanks, and 3) model geographic regions in need of further taxonomic exploration.

## 2. METHODS

### 2.1 Taxon sampling

IUCN Red List shapefiles were imported into QGIS for mapping All currently described New World monkey (Platyrrhini) distributions (https://www.iucnredlist.org/), representing 161 species and 34 subspecies. Distinctions were made between points with GPS data, general geographic descriptions, and those based on IUCN shapefiles (Supplemental Table S1).

### 2.2 Riverine analyses

Geographic boundaries (i.e., rivers, mountains, plateaus, etc.) of all adjacent species and subspecies were measured using Google satellite maps. Annual river flow rates (m3/s) and river width (km) were used from Ayres & Clutton-Brock (1992) and Fordham et al. (2020), when available (Supplemental Table S2). We also used the Global Runoff Data Centre database to ascertain river measurements from streamflow-monitoring stations (GRDC, Germany). River width was measured in the midpoint of species boundaries from available satellite imagery using a previously published methodology (Fordham et al. 2020).

### 2.3 Molecular phylogenetics

All molecular data and associated sampling locations used in the main river barrier analysis were obtained from NCBI (https://www.ncbi.nlm.nih.gov/). Partial mitochondrial sequences (i.e., cytochrome b and cytochrome oxidase subunit II) were trimmed to the smallest consensus size for both overall phylogenetic analysis and analysis between adjacent or sympatric taxonomic groups. A minimum of 200bp sequence and a maximum of 1104bp were available. The two mitochondrial genes were not concatenated but analyzed separately to independently verify results since sequence data was not necessarily available from the same individuals and for the same taxonomic groups. Each outgroup was chosen from the sister group, Catarrhini, using the closest available sequences for both cytb (NC_056330.1) and COII (NC_006901.1). We used MAFFT (version 7.407_1) for sequence alignment and curated with trimAI (version 1.41) (Capella-Gutierrez et al., 2009). The phylogenetic tree was built using distance-based inference of phylogenetic trees with combined PhyML + SMS programs (version 1.8.1_1). Molecular similarity was tested for normality using the Shapiro-Wilk W test, while the Mann-Whitney U test was used to compare nonparametric data between species and subspecies data for both molecular markers. Kruskal-Wallis nonparametric analysis of variance was used to compare molecular similarity values in taxonomic groups constrained by rivers, mountains, and those with no observable geographical barriers for both molecular markers. River characteristics (i.e., width and flow rate) were compared to the molecular similarity of adjacent taxa on opposing riverbanks using Spearman’s nonparametric rank coefficient.

## 3. RESULTS

### 3.1 Geographical barriers

Rivers were found at the interface of species boundaries in 73.9% (198/268) of analyzed cases, while mountains separated currently described species in 5.2% of cases (14/268). No observable geographical barriers were seen at the interface of 18.7% of species ranges. The majority (78.6%) of shared taxonomic boundaries (species and subspecies) were separated by geographical barriers such as rivers or mountains (236/300); however, several currently described genera had species interfaces with no geological barrier: *Lagothrix* (1/1 species), *Mico* (6/22 species), *Callicebus* (4/5 species), *Plecturocebus* (12/43 species), *Cebus* (6/18 species), and *Sapajus* (5/12 species). At subspecies boundaries, geographical barriers were present in 75% of cases (24/32 species).

The average width of rivers at all adjacent species boundaries was 1.0 km while subspecies had an average river width of 0.12 km. Subspecies were separated by rivers no larger than 0.22 km in width (Table 1). In general, wider rivers were associated with an increased molecular dissimilarity. The greatest average molecular dissimilarity (90%: cytb) was found on the Negro River, with an average width of 1.95 km at adjacent taxa boundaries. The rivers Branco, Jiparana, and Amazon all had average molecular similarity equal or less than 95% (cytb) while only the Jurua River had 95% similarity or less (COII). Several rivers were found separating multiple Platyrrhini taxa: the Negro River (1.95 km width; N= 2 genera, 2 species); Amazon (1.11 km; N=4,10), Madeira (1.2 km; N=1,3); Japurá (0.71 km; N=2,3); Tapajós (2.23 km; N=4,4), Marañón (0.61 km; N=3,3); Huallaga (0.58 km; N=2, 2); São Francisco (1.02 km; N=2,3); Purus (0.3 km; N=2,2); Juruá (0.28 km; 3,3), Jequitinhonha (0.23 km; N=2,2).

**Table 1:**
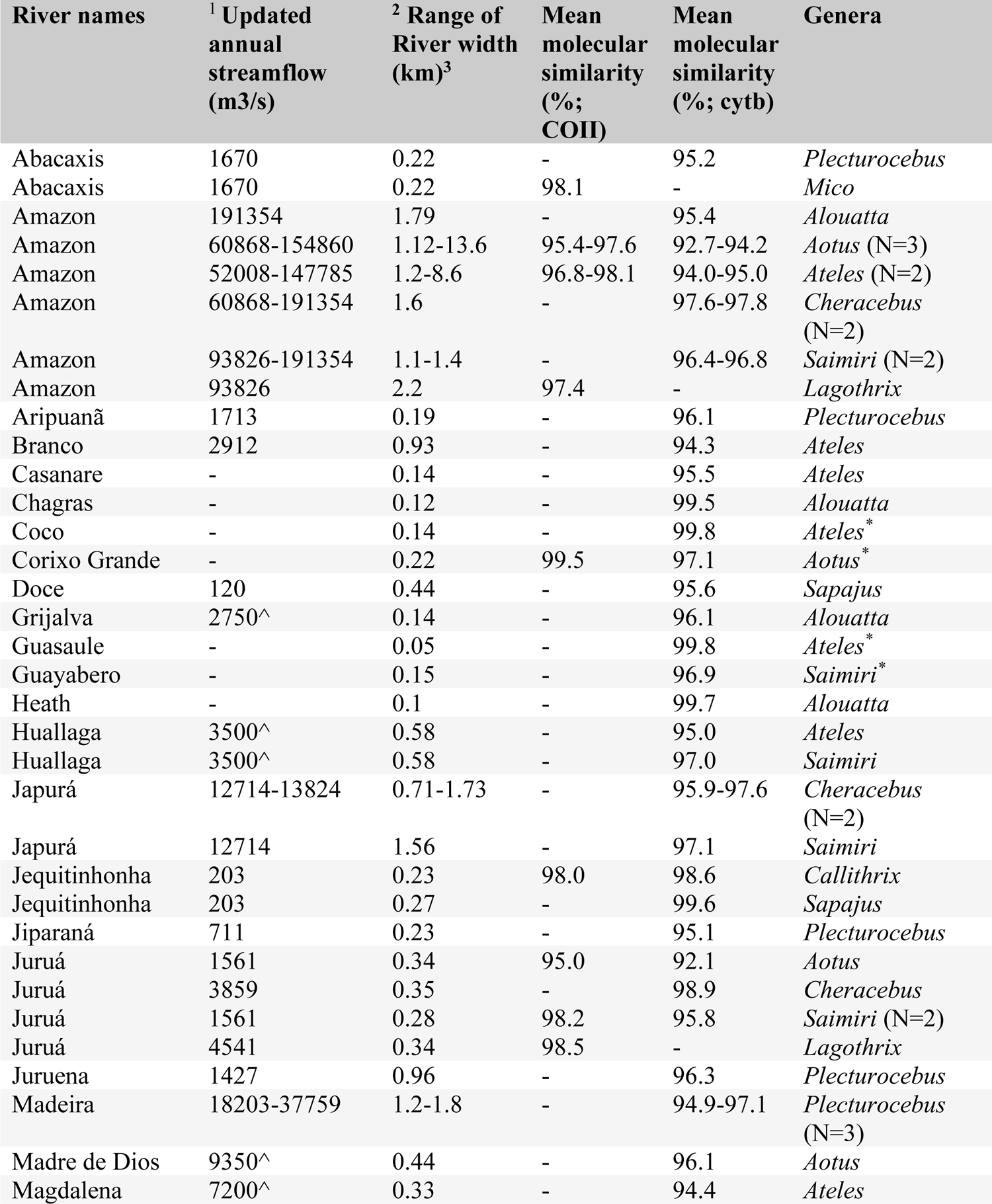
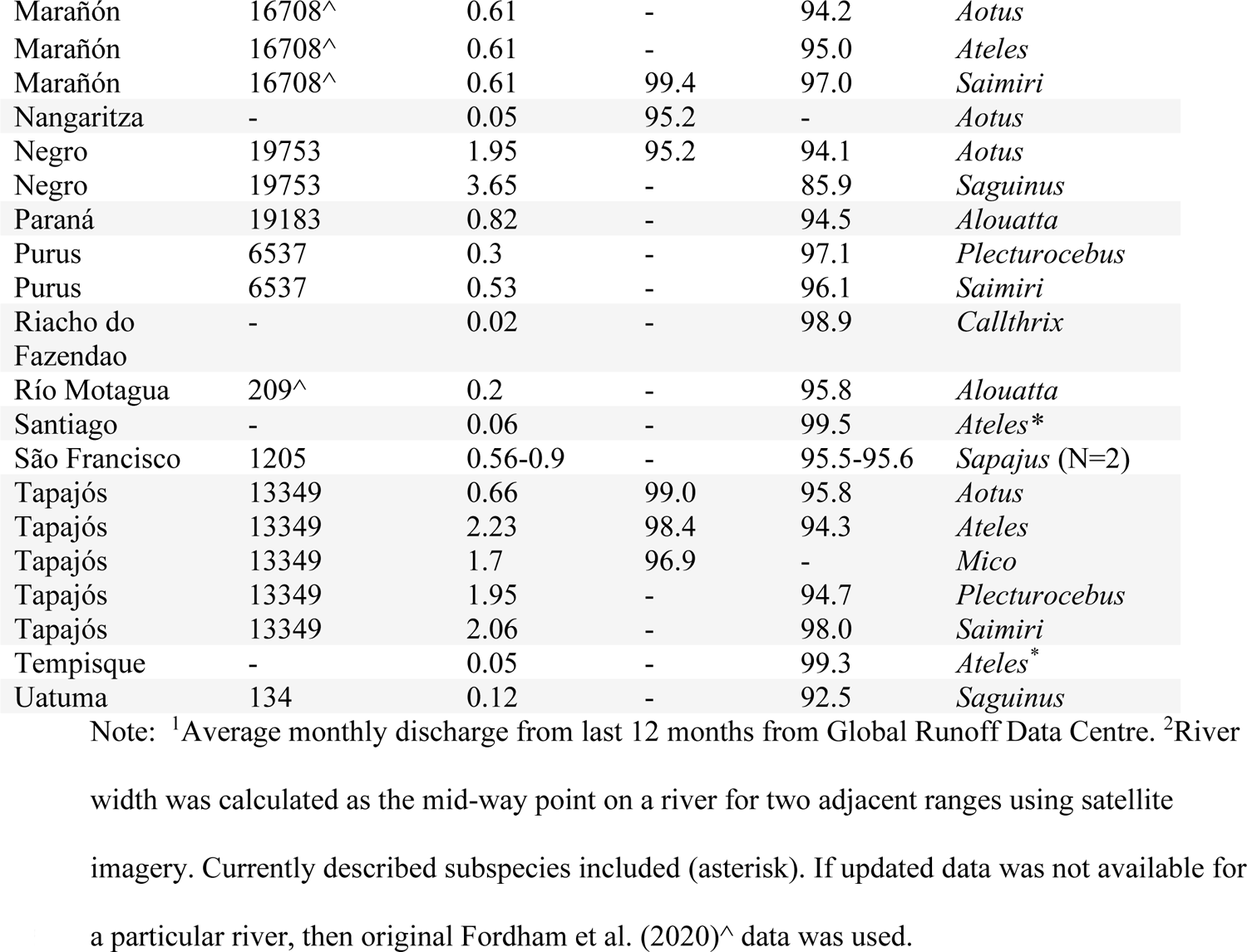
River characteristics and molecular similarity for adjacent taxonomic groups on opposing riverbanks.

### 3.2 River width and molecular dissimilarity

River width was positively correlated with the molecular dissimilarity of adjacent taxa on opposing riverbanks using both cytb (non-parametric Spearman’s Rank Order Correlation: N=59; r=0-.46; p=0.00; Figure 1a), and a smaller set of COII sequences which were not significant (non-parametric Spearman’s Rank Order: N=18; r =-0.23; p=0.36; Figure 1c). Annual streamflow (m3/s) was not significantly associated with molecular dissimilarity in adjacent taxa on opposite riverbanks for neither cytb (Spearman’s Rank Order Correlation: N=46; r=0-.21; p=0.16) nor COII (N=16; r=-0.30; p=0.26; Figure 1b,c), though there was a generally positive relationship. Several data points in all four analyses fell above and below the 95% confidence interval. However, when data from adjacent taxa with no river barrier were included along with unsupported samples (i.e., 100% molecular similarity between adjacent taxa), there was a similar – but significant - trend for both cytb (N=92; r=-0.37; p=0.00) and COII (N=31; r=-0.42; p=0.02).

**Figure 1.**
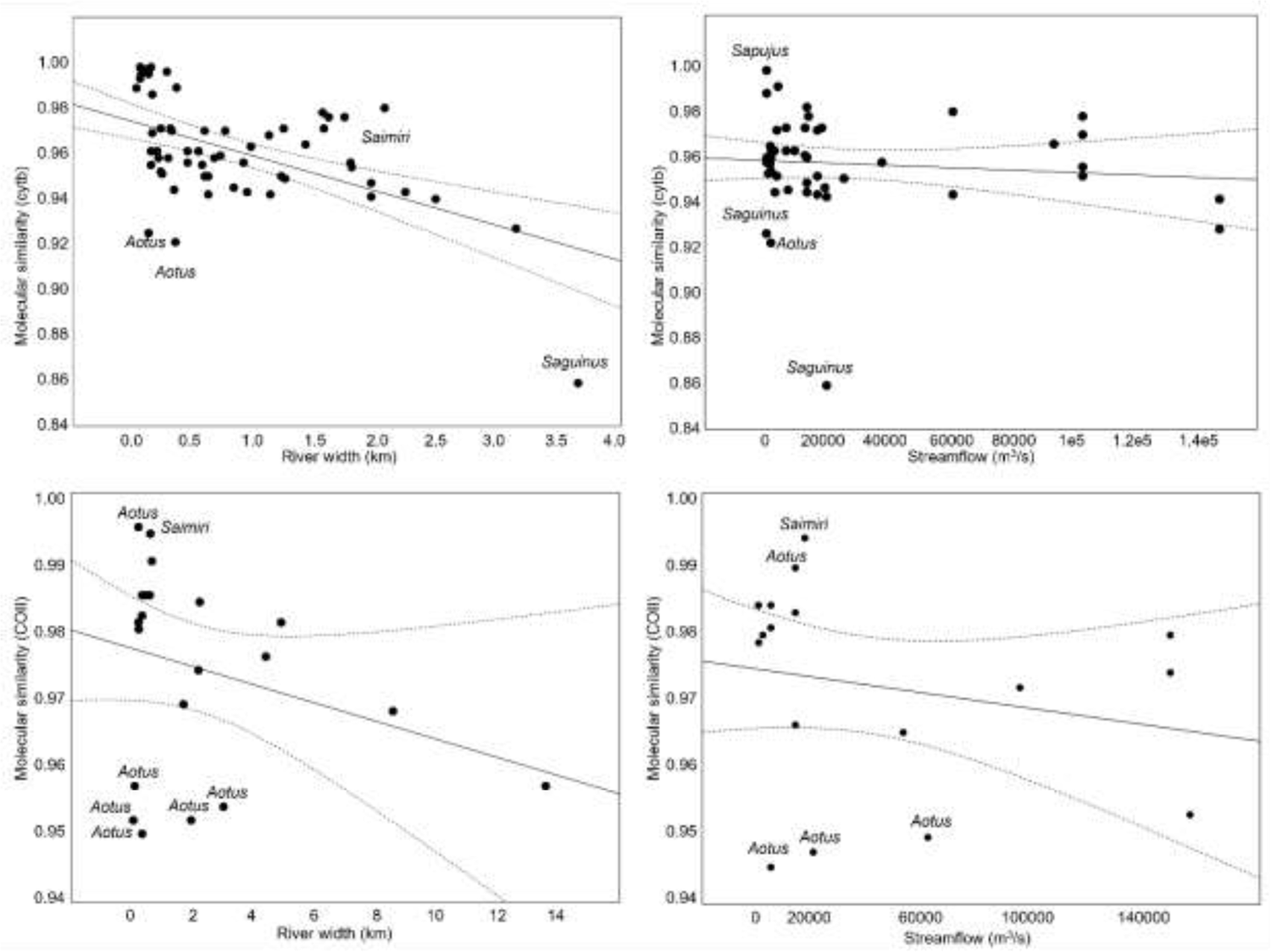
Comparison of molecular similarity of adjacent taxa on opposing sides of river coupled with river width (km). Upper figures (a,b) represent analysis using partial cytochrome b (cytb) sequence and lower figures (c,d) use available cytochrome oxidase subunit II (COII) data. Ninety-five percent confidence interval represented by dashed lines.

Molecular similarity within each genus was negatively correlated with river width in all cases except for *Saimiri* (cytb; Spearman R = 0.53, N=10, p=0.11; Figure 2). However, only *Alouatta* (cytb, Spearman R = −0.94; N=6; p<0.05; Figure 2c) and *Ateles* (cytb, Spearman R = − 0.87; N=12; p=0.00; Figure 2b) exhibited a significant intra-genus negative relationship between wider river and increased molecular dissimilarity while *Callithrix* and *Saguinus* both had insufficient data for both molecular markers. *Saimiri* exhibited a similar trend for COII sequence.

**Figure 2.**
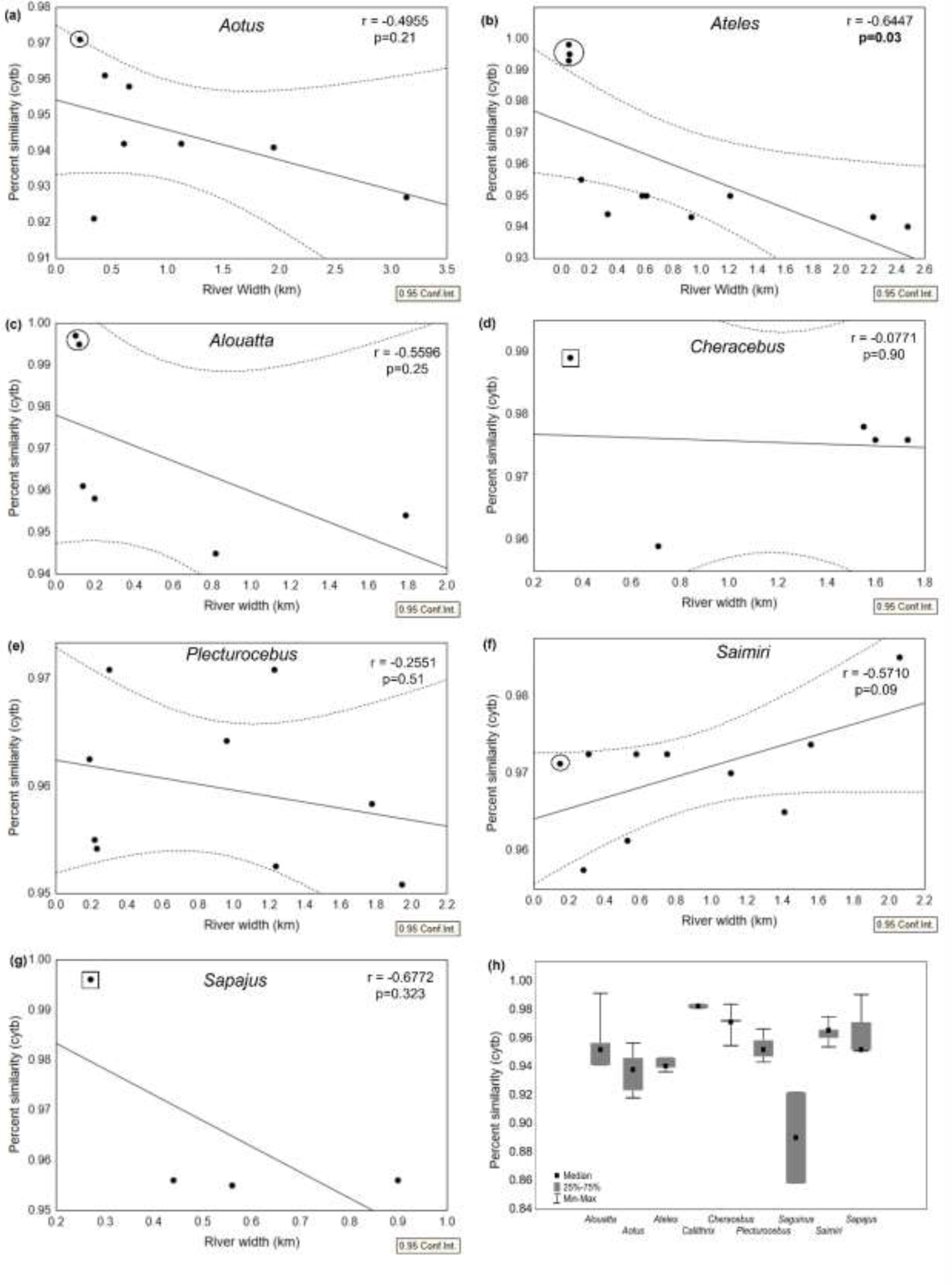
(a-g) Association of river width (km) with cytb sequence similarity (y-axis) of adjacent species on opposing sides of barrier for each Platyrrhini genus with available data. *Saguinus* only included two data points and was not included. (h) Descriptive statistics for cytb sequence similarity data from (a-g), depicting those taxa with far lower median values compared to others (e.g. *Saguinus* and *Aotus*) and those taxa with upper ranges indicative of subspecies or possibly misclassified taxa (e.g., *Callithrix*, *Cheracebus*, *Saimiri*, and *Sapajus*). Dotted lines represent linear trendlines and include the correlation coefficient (r). Curved outer bands represent 95% confidence interval. Encircled points represent presently described subspecies while squares depict taxa presently described as species that have similar phylogenetic similarities consistent with subspecies classification.

Several genera exhibited greater molecular divergence (i.e., lower similarity) on opposing riverbanks than others. *Saguinus* exhibited a far lower average molecular similarity range of values (Figure 2h: cytb mean: 89.2%; range: 85.9-92.5%) compared to most other genera though only statistically different from *Cherecebus* (cytb mean: 97.6%; range: 95.9-98.9%; p<0.05). The range of *Aotus* molecular similarity values from taxa on opposing riverbanks (Figure 2a: cytb mean: 94.0%; range: 92.1-96.1%) was also significantly lower than *Cheracebus* (Figure 2d: cytb mean: 97.6%; range: 95.9-98.9%; p<0.05) and *Saimiri* (Figure 2f: cytb mean: 96.9%; range: 95.8-98.0%; p<0.05). COII data set was comparatively smaller and no statistically significant results were found between genera.

### 3.3 Molecular phylogenetic trees

Seventy-five percent (Figure 3: cytb: 66/88) and 90.5% (COII: 19/21) of phylogenetic trees comparing all sequences within two adjacent taxonomic groups were supported. Unsupported molecular differentiation between adjacent taxonomic groups separated by rivers was 24.2% (cytb: N=66) and 10.5% (COII: N=19) of cases compared to adjacent taxonomic groups with no observable geographical barrier (cytb: 55.6%; N=18 and COII: 11.1%; N=9). All taxonomic groups separated by mountains were supported molecularly for both mitochondrial genes.

**Figure 3.**
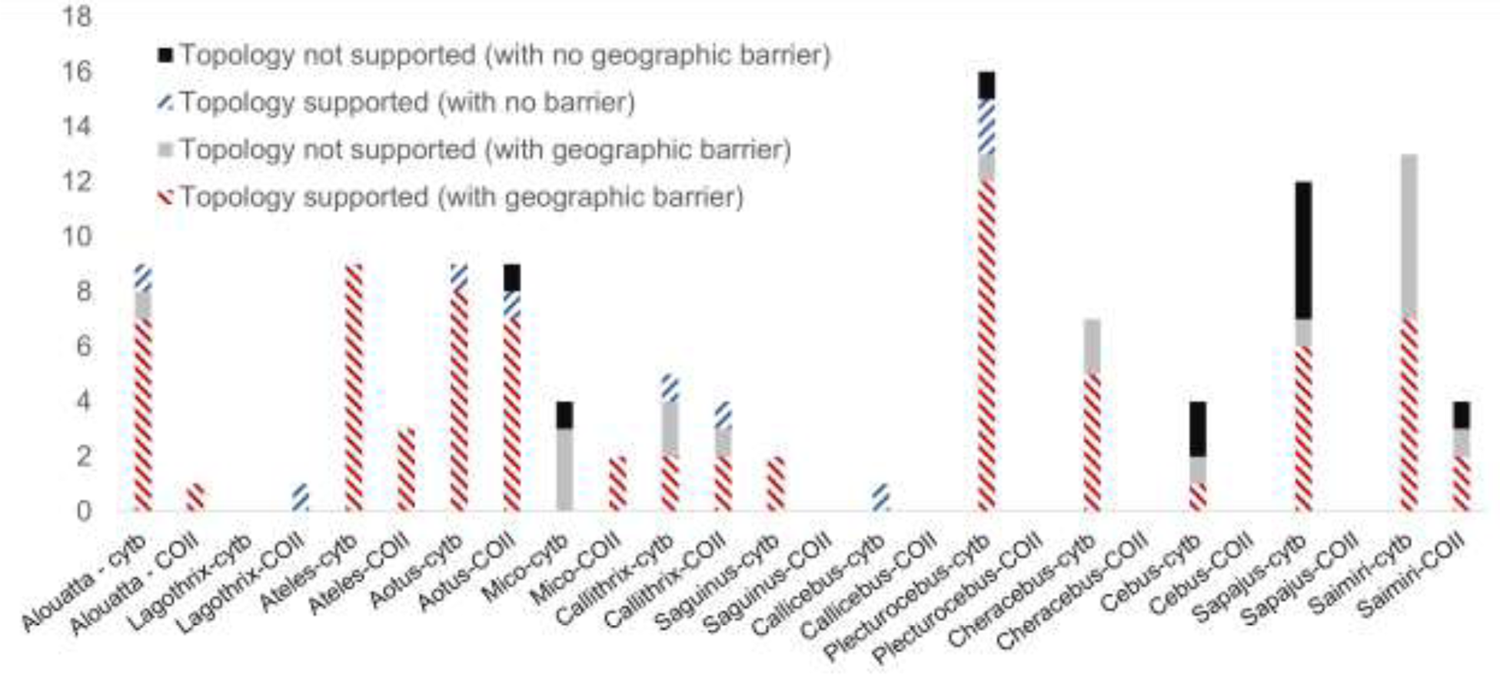
Phylogenetic support for adjacent species with geographic barriers versus those without any river or mountain barrier. Does not include subspecies analysis. Molecular data was not available for *Brachyteles*, *Cebuella*, *Leontopithecus*, *Leontocebus*, and *Pithecia*.

Several individual river barriers had statistically higher levels of molecular dissimilarity than anticipated. *Saguinus inustus* and *S. midas* only shared 85.9% molecular similarity (cytb) despite being adjacent species, though they were separated by one of the largest rivers in the Amazon Basin, the Negro River. Two other barriers had higher molecular dissimilarity than expected based on narrow river width: *Aotus nancymaae* and *A. nigriceps* were 92.1% molecularly similar despite being separated by the Juruá River which is only 0.34 km wide, and *Saguinus midas* and *S. bicolor* were only 92.5% similar despite being separated by the Uatuma river with a width of only 0.12 km. Conversely, unexpectedly high molecular dissimilarity on opposing riverbanks was found for *Saimiri ustus* and *S. collinsi*. For example, the Tapajós River was over 2 km wide at a minimum, and yet these two species had 98% molecular similarity.

Well-resolved phylogenetic trees were found in 71.7% of cases using partial cytb sequences and 87.0% of cases using partial COII sequences – irrespective of whether physical barriers were present or not. For all adjacent species comparisons, mean molecular similarities were 95.6% (cytb: N=51; Interquartile range 50%: 94.5-97.0%) and 97.2% (COII: N=16; Interquartile range 50%: 95.6-98.2); while subspecies level differences were 98.5% (cytb: N=8; Median 99.4%; Interquartile range 50%: 97.0-99.65%) and 99% (COII: N=2; Interquartile range 50%: 98.5-99.5%). The difference in mean molecular similarity between species versus subspecies similarity was significant using cytb (Mann-Whitney: p<0.001). Mean COII molecular similarity between species and subspecies was not significant (Mann-Whitey: p=0.06).

No statistical difference was found between molecular similarity across all taxonomic groups separated by rivers and mountains versus those with no discernable geographical barrier (Kruskal-Wallis (cytb: ꭓ^2^=0.39, p=0.82, df=2; COII: ꭓ^2^=3.67, p=0.16, df=2). An average molecular similarity was found of 95.5% (cytb: N=53) and 96.9% (COII: N=17) when river boundaries were present (range=89.2-98.8%), 96.3% (cytb) and 94.3% (COII) when mountain barriers were present (range=94.3-98.5%), and 95.6% (cytb) and 96.5% (COII) where no observable geological barrier was present (range=93.6-98.2%). Among subspecies, molecular similarity among adjacent taxa was on average 98.3% similar (cytb) and 96.7% (COII) on opposing geographical barriers, while adjacent taxa where no observable geographical barrier existed were 96.2% (cytb; N=2) and 99.0% (COII; N=1) similar. No subspecies were found to be separated by mountains.

### 3.5 Suspect taxonomy

Species delimitation was not supported in taxa on opposing river barriers in 25.8% (cytb) and 10.5% (COII) of analyzed cases (Table 2). In cases with no geographical barrier between presently described adjacent taxa species delimitation was not supported in 62.5% (cytb) and 11.1% (COII) of cases. Unsupported phylogenetic relationships were found in only two instances where mountains separated taxonomic groups. Adjacent taxa that were not supported phylogenetically for either molecular marker were *Aotus azarai infulatus* and *A. azarai boliviensis* (now a proposed species by Martins-Junior 2022), *Callithrix geoffroyi* and *C. penicillata*, *Saimiri boliviensis boliviensis* and *S. b. peruviensis*, and *Saimiri oerstedii oerstedii* and *S. o citrinellus.* Numerous other adjacent taxa were supported phylogenetically only at a single molecular marker, while for many other relationships data wasn’t available for both species involved.

**Table 2:** Evidence of unsupported topologies for at least one molecular marker (or both) between presently described adjacent taxonomic groups. cytb and COII values represent percent sequence similarity. All *Saguinus* relationships were supported.

| Genus | Species sharing adjacent ranges | cytb | COII | River Width (km) | Fordham 2020 Methodology* |
| --- | --- | --- | --- | --- | --- |
| *Alouatta* | *A. macconnelli* and *A. seniculus* | Unsupported | No data | 1.95 | 19753 |
| *Aotus* | *A. azarai infulatus* and *A. a. boliviensis* | Unsupported | Unsupported | 0 | 0 |
|  | *A. azarai azarai* and *A. a. boliviensis* | 97.7% | Mixed | 0 | 0 |
| *Callithrix* | *C. geoffroyi* and *C. penicillata* | Unsupported | Unsupported | 0.07 | No data |
|  | *C. penicillata* and *C. aurita* | Unsupported | 94.3% | 0 | NA |
| *Cheracebus* | *C. purinus* and *C. torquatus* | Unsupported | No data | 1.23 | No data |
|  | *C. lugens* and *C. torquatus* | Unsupported | No data | 0.06 | No data |
| *Cebus* | *C. albifrons* and *C. castaneus* | Unsupported | No data | 0.93 | 2912 |
|  | *C. albifrons* and *C. olivaceus* | Unsupported | No data | 0 | 0 |
|  | *C. olivaceus* and *C. castaneus* | Unsupported | No data | 0 | 0 |
| *Lagothrix* | *L. lagotricha poeppigii* and *L. l. tschudii* | No data | Unsupported | 0.67 | No data |
|  | *L. lagotricha lugens* and *L. l. lagotricha* | No data | Unsupported | 0 | 0 |
|  | *L. lagotricha lagotricha* and *L. l. poeppigii* | No data | Unsupported | 0 | 0 |
| *Mico* | *Mico humeralifer* and *M. argentatus* | Unsupported | 96.9% | 0.96 | 13349 |
|  | *Mico mauesi* and *M. humeralifer* | Unsupported | No data | 0.18 | No data |
|  | *Mico mauesi* and *M. chrysoleucos* | Unsupported | No data | 0.48 | No data |
|  | *Mico mauesi* and *M. melanurus* | Unsupported | No data | 0 | 0 |
| *Plecturocebus* | *Plecturocebus discolor* and *P. cupreus* | Unsupported | No data | 0.67 | No data |
|  | *Plecturocebus moloch* and<br>*P. vieirai* | Unsupported | No data | 0 | 0 |
| *Saimiri* | *Saimiri boliviensis*<br>*boliviensis* and *S. b.*<br>*peruviensis* | Unsupported | Unsupported | 0 | 0 |
|  | *Saimiri cassiquiarensis*<br>*cassiquiarensis* and *S.*<br>*sciureus* | Unsupported | No data | 2.35 | 19753 |
|  | *Saimiri cassiquiarensis*<br>*cassiquiarensis* and *S.*<br>*ustus* | Unsupported | No data | 2.73 | 191354 |
|  | *Saimiri cassiquiarensis*<br>*macrodon* and *S.*<br>*vanzolinii* | Unsupported | No data | 0.76 | No data |
|  | *Saimiri ustus* and *S.*<br>*sciureus* | Unsupported | No data | 2.5 | 191354 |
|  | *Saimiri sciureus* and *S.*<br>*collinsi* | Unsupported | No data | 4.16 | 191354 |
|  | *Saimiri oerstedii*<br>*oerstedii* and *S. o*<br>*citrinellus* | Unsupported | Unsupported | 0.09 | No data |
|  | *Saimiri ustus* and *S.*<br>*vanzolinii* | Unsupported | No data | 1.2 | No data |
|  | *Saimiri cassiquiarensis*<br>*macrodon* and *S. c.*<br>*albigena* | Unsupported | No data | 0 | NA |
| *Sapajus* | *Sapajus nigritus* and<br>*S. cay* | Unsupported | No data | 0.82 | 19183 |
|  | *Sapajus apella apella*<br>and<br>*S. libidinosus* | Unsupported | No data | 0 | 0 |
|  | *Sapajus apella apella*<br>and<br>*S. cay* | Unsupported | No data | 0 | 0 |
|  | *Sapajus flavius* and<br>*S. libidinosus* | Unsupported | No data | 0 | 0 |
|  | *Sapajus nigritus* and<br>*S. libidinosus* | Unsupported | No data | 0 | 0 |
|  | *Sapajus libidinosus* and<br>*S. cay* | Unsupported | No data | 0 | 0 |

\*Annual streamflow (m3/s) using average monthly discharge from last 12 months.

Of note, *Lagothrix* had three unsupported adjacent taxa (COII) out of seven interspecies borders. No cytb data was available for *Lagothrix*. All four bordering *Mico* adjacent species were phylogenetically unsupported (cytb only), though one of these cases was supported using COII. Three of these mixed results were at river boundaries and one with no geographical barrier.

*Saimiri* had nine unsupported phylogenetic cases. No shared *Saimiri* species borders were supported molecularly using cytb, though two adjacent taxa were supported using COII data. Six bordering *Saimiri* had river barriers while one subspecies case had a mountain range, and another had no geographical barrier. Of four bordering *Cebus* adjacent species with available data (only cytb), only one barrier was supported (mountains), while two had no geographical barrier and another was separated by the Branco River (0.6km width). Finally, six bordering *Sapajus* species were not supported phylogenetically (only cytb available) for which five had no geographical barrier.

## 4. DISCUSSION

The findings of this study demonstrate the significant role of rivers as primary geographical barriers that separate species and subspecies of Platyrrhini primates. Rivers were frequently observed at species boundaries, while mountains played a lesser role in species separation. Specifically, we identified 35 rivers across Central and South America that effectively restricted gene flow between adjacent pairs of New World monkey species. Larger rivers such as the Amazon, Japura, Madeira, and Tapajós were found to separate three or more genera of primates and exhibited greater molecular dissimilarity on opposing riverbanks. Smaller rivers like the Riacho do Fazendao and the Nangaritza also showed associations with divergent molecular signatures in adjacent species. These results highlight the crucial role of rivers in shaping the genetic differentiation and diversity of Platyrrhini primates.

Our findings are consistent with previous research conducted by Ayres & Clutton-Brock (1992) and Fordham et al. (2019), which demonstrated a negative correlation between river size and species composition similarity on opposite riverbanks. Similar patterns have been observed in other taxa, such as plants (Nazareno et al., 2019), amphibians (Moraes et al., 2016; Figueiredo-Vázquez et al., 2021), and birds (e.g., Capparella, 1987; Hayes & Sewlal, 2004; Fernandes et al., 2012; Ribas et al., 2012; Pomara et al., 2014; Naka & Brumfield, 2018; Kopuchian et al., 2020), where larger river widths are associated with higher levels of genetic differentiation. Additionally, when comparing molecular similarity between taxa on opposing riverbanks, we observed varying levels of molecular divergence among genera. Saguinus exhibited significantly lower average molecular similarity compared to most other genera, indicating higher genetic differentiation. Aotus also showed lower molecular similarity values compared to *Cheracebus* and *Saimiri*. Furthermore, our study revealed that certain genera like *Lagothrix*, *Mico*, *Callicebus*, *Plecturocebus*, *Cebus*, and *Sapajus* exhibited species interfaces despite the absence of geological barriers, suggesting that other factors, such as ecological or behavioral differences, contribute to maintaining species boundaries in these cases.

Our findings revealed a positive correlation between river width and molecular dissimilarity across all genera, indicating that wider rivers act as stronger barriers to gene flow. The width of rivers at species boundaries varied, with an average width of 1.0 km for all adjacent species boundaries and 0.12 km for subspecies boundaries. Notably, the Negro River, with the widest average width (1.95 km) at adjacent taxa boundaries, exhibited the greatest average molecular dissimilarity. Other rivers such as the Branco, Jiparana, and Amazon also showed high molecular dissimilarity, suggesting their role as barriers to gene flow. Although the association between annual streamflow and molecular dissimilarity was not significant, a general positive pattern was observed when considering unsupported phylogenetic results and adjacent species similarity with no river barrier.

We further investigated the relationship between river width and molecular dissimilarity within each genus. Except for *Saimiri*, all other genera showed a negative correlation between river width and molecular similarity, indicating that wider rivers are associated with increased molecular dissimilarity within genera. However, presumably due to a small number of species, all but *Ateles* was insignificant within genera. Notably, *Alouatta* and *Ateles* exhibited significant intra-genus negative relationships, indicating a strong impact of wider rivers on genetic differentiation within these genera – despite their high level of mobility. Some specific cases highlighted unexpected levels of molecular dissimilarity between adjacent taxa separated by geographical barriers. For example, *Saguinus inustus* and *S. midas* exhibited particularly low molecular similarity (e.g., cytb: 85.9% similarity) even for a relatively wide river (i.e., Negro River: 3.65 km width). A limited number of data points were available for *Callithrix* and *Saguinus*, preventing definitive conclusions. These findings underscore the genus-specific differences in the influence of river width on genetic differentiation.

The ability of Platyrrhini taxa to traverse rivers or occupy different ecological niches associated with dispersal such as canopy and river edge species reported in birds (Burney & Brumfield, 2009), may explain why some species can cross larger rivers while others cannot. This alone may not be sufficient to explain why some species are capable of crossing larger rivers. Body size alone does not appear to be a determining factor, as relatively smaller species like capuchins (i.e., *Sapajus* and *Cebus*) are equally capable of crossing larger rivers compared to larger species like *Ateles*. For example, the relatively smaller, lower canopy capuchin species are not restricted by larger rivers while the large and very mobile *Ateles* is often found in the upper canopy but is limited by the Tapajos and Amazon Rivers (Lima et al., 2018). *Cheracebus* (mean male weight: 1.5 kg) had the highest molecular similarity when controlling for river width (cytb: 97.7%) despite being far smaller than *Alouatta* (mean male weight: 9.8 kg) or *Ateles* (mean male weight: 11 kg). However, further research is needed to elucidate the specific mechanisms behind this phenomenon.

While hybridization and introgression are common along species edges of some Platyrrhini genera (e.g., Bicca-Marques et al., 2008; Mercês et al., 2015; Carneiro et al., 2016), many currently described adjacent species were not supported molecularly. Nevertheless, adjacent species were more phylogenetically similar when no geographical barrier was evident. This suggests the importance of incorporating geographic barriers into taxonomic descriptions. However, it is important to note that the two molecular markers used in this study may not be sufficient for distinguishing adjacent species. The intention was not to describe new species solely on two partial sequences, but as a quantitative measure of divergence. Further evidence from multiple disciplines should therefore be considered.

*Aotus* and *Saguinus* were far more dissimilar on opposing riverbanks than expected based solely on river size. Small rivers were sufficient to restrict dispersal in both genera, leading to higher molecular dissimilarity than expected based on overall relationship across all other New World monkey adjacent species despite their proficient arboreal nature. As an example, the Jurua River is only 0.34 km in width and was a barrier between adjacent species of *Aotus* (cytb and COII), C*heracebus* (cytb and COII), *Saimiri* (cytb and COII), and *Lagothrix* (COII). However, *Aotus* showed the lowest molecular similarity between adjacent species on opposing riverbanks (cytb: 92.1% and 95.0%: COII). *Lagothrix* and *Saimiri* had the highest molecular similarity on opposing riverbanks (COII: 98.5% and 98.2%, respectively). The potential existence of new taxonomic diversity within several genera, including *Aotus*, has important conservation implications. The range of *Aotus nigriceps*, for example, encompasses a vast area in the Amazon basin that is currently facing significant deforestation and wildfires (Helenbrook & Valdez 2021). The description of additional taxonomic diversity can provide important information for conservation planning and maximize protections for neotropical primate species and intraspecies diversity in these highly threatened habitats.

This study has several limitations that should be considered. Comparisons within genera were constrained by limited sample sizes, and a significant number of Platyrrhini species lacked available data for molecular analysis. In some instances, sequence data was accessible, but associated geographic coordinates were missing. Nonetheless, it may be feasible to deduce molecular divergence from geographical barriers, which could aid in identifying potential areas that might harbor new distinct taxa by incorporating biogeographical evidence. However, it is crucial to recognize that predictions of molecular divergence inferred solely from biogeography do not suffice to describe new taxa, and further research is necessary. The integration of multiple lines of evidence, including morphology, paleontology, behavior, and ecology, is crucial for a comprehensive understanding of species boundaries and genetic differentiation within Platyrrhini primates (Costa-Araujo et al., 2021).

Regarding the relationship between geographic barriers and speciation, there are significant questions regarding the history of riverine formation in the neotropics and their contribution to species richness and maintenance of contemporary species (e.g., Naka and Brumfield 2018; Méndez-Camacho et al. 2021). Estimates of river formation in the Amazon basin, ranging from 2-12 million years contribute to uncertainty (Rossetti et al., 2005; Campbell et al., 2006; Figueiredo et al., 2009; Hoorn et al. 2010; Latrubesse et al. 2010; Latrubesse et al. 2010; Hoorn 2010; Ribas et al. 2012; Méndez-Camacho et al. 2021). While the formation of a river barrier preceding the origin of taxonomic divergence is not sufficient evidence to infer causation, additional evidence of endemism and other biogeographical patterns can provide further insights (Silva et al. 2002; Borges and Silva 2012).

Nonetheless, it is important to consider evidence of endemism that does not always align with river boundaries, as demonstrated by studies conducted by Hall and Harvey (2002) and Oliveira et al. (2017). Discrepancies may in part be explained by fluctuating river characteristics both due to sediment deposition in upper headwaters which allow dispersal, rearrangement of river networks (Ruokolainen et al. 2019), or more complex evolutionary processes. Nonetheless, our results quantitatively support the River Barrier Hypothesis despite these potentially confounding processes. Rather this study points to the importance of geological barriers in restricting dispersal, and how it could be used as a tool to predict additional taxonomic boundaries.

## 5. CONCLUSIONS

This study emphasizes the crucial role of rivers as primary geographical barriers that shape the genetic differentiation and diversity of Platyrrhini primates. In light of the alarming deforestation rates in New World monkey habitats, driven by agricultural expansion and fires in the Amazon rainforest (Cardil et al., 2020; Helenbrook & Valdez, 2021), urgent action is needed to protect these highly vulnerable primate species. Adopting a systematic approach that integrates statistical biogeography, molecular phylogenetics, and multiple lines of evidence is essential for better understanding species boundaries, genetic differentiation, and guiding targeted conservation efforts. Furthermore, the potential existence of new taxonomic diversity within Platyrrhini genera underscores the importance of comprehensive species descriptions and conservation planning to safeguard neotropical primate diversity.

## CONFLICT OF INTEREST

The author acknowledges no potential conflict of interest.

## DATA AVAILABILITY

The raw data supporting the conclusions of this article are included in the article or as Supplementary Material (Supplemental Table T1 and T2).

## BIOSKETCH

William Helenbrook is the Research Director for the Tropical Conservation Fund (TCF) and Adjunct Assistant Professor with SUNY College of Environmental Science and Forestry. He collaborates with Peruvian and Brazilian NGOs on research related to neotropical primate conservation biology, phylogenetics, conservation genomics, and disease ecology. He also engages local communities and partners to protect tropical forests through environmental education, ecotourism, creation of biodiversity offsets, and applied research.

## Notes

### Competing Interest Statement

The authors have declared no competing interest.

## REFERENCES

Aleixo, A. (2004). Historical diversification of a terra-firme forest bird superspecies: a phylogeographic perspective on the role of different hypotheses of Amazonian diversification. Evolution 58(6), 1303–1317.

Alfaro, J. W. L., Boubli, J. P., Paim, F. P., Ribas, C. C., da Silva, M. N. F., … Farias, I. P. (2015). Biogeography of squirrel monkeys (genus *Saimiri*): South-central Amazon origin and rapid pan-Amazonian diversification of a lowland primate. Molecular Phylogenetics and Evolution 82, 436–454.

Ayres, J. M., & Clutton-Brock, T. H. (1992). River boundaries and species range size in Amazonian primates. The American Naturalist 140(3), 531–537.

Beaudrot, L. H., & Marshall, A. J. (2011). Primate communities are structured more by dispersal limitation than by niches. Journal of Animal Ecology 80(2), 332–341.

Bicca-Marques, J. C., Prates, H. M., de Aguiar, F. R. C., & Jones, C. B. (2008) Survey of Alouatta caraya, the black-and-gold howler monkey, and Alouatta guariba clamitans, the brown howler monkey, in a contact zone, State of Rio Grande do Sul, Brazil: evidence for hybridization. Primates 49(4), 246–252.

Borges, S. H. & Da Silva, J. M. C. (2012). A new area of endemism for Amazonian birds in the Rio Negro Basin. Wilson Journal of Ornithology 124, 15–23.

Boubli, J. P., Ribas, C., Alfaro, J. W. L., Alfaro, M. E., da Silva, M. N. F., Pinho, G. M., & Farias, I. P. (2015). Spatial and temporal patterns of diversification on the Amazon: A test of the riverine hypothesis for all diurnal primates of Rio Negro and Rio Branco in Brazil. Molecular Phylogenetics and Evolution 82, 400–412.

Boubli, J. P., Byrne, H., da Silva, M. N., Silva-Júnior, J., Araujo, R. C., … Farias, I. P. (2019). On a new species of titi monkey (Primates: Plecturocebus Byrne et al., 2016), from Alta Floresta, southern Amazon, Brazil. Molecular Phylogenetics and Evolution 132, 117–137.

Burney, C. W., & Brumfield, R. T. (2009). Ecology predicts levels of genetic differentiation in Neotropical birds. The American Naturalist 174(3), 358–368.

Bush, M. B. (1994). Amazonian speciation: a necessarily complex model. Journal of Biogeography 5–17.

Bush, M. B., & Oliveira, P. E. D. (2006). The rise and fall of the Refugial Hypothesis of Amazonian speciation: a paleoecological perspective. Biota Neotropica 6.

Capparella, A. P. (1987). Effects of riverine barriers on genetic differentiation of Amazonian forest undergrowth birds (Peru). Doctoral dissertation, Louisiana State University and Agricultural & Mechanical College).

Capella-Gutiérrez, S., Silla-Martínez, J. M., & Gabaldón, T. (2009). trimAl: a tool for automated alignment trimming in large-scale phylogenetic analyses. Bioinformatics 25(15), 1972–1973.

Cardil, A., De-Miguel, S., Silva, C. A., Reich, P. B., Calkin, D., Brancalion, P. H., … & Liang, J. (2020). Recent deforestation drove the spike in Amazonian fires. Environmental Research Letters 15(12), 121003.

Carneiro, J., Rodrigues-Filho, L., Schneider, H., & Sampaio, I. (2016). Molecular data highlight hybridization in squirrel monkeys (*Saimiri*, Cebidae). Genetics and Molecular Biology 39, 539–546.

Chapman, F. M. (1926). The distribution of bird-life in in Ecuador: A contribution to a study of the origin of Andean bird-life. Bulletin American Museum of Natural History 55, 1– 784.

Chiou, K. L., Pozzi, L., Alfaro, J. W. L., & Di Fiore, A. (2011). Pleistocene diversification of living squirrel monkeys (*Saimiri* spp.) inferred from complete mitochondrial genome sequences. Molecular Phylogenetics and Evolution 59(3), 736–745.

Costa-Araújo, R., Boubli, J. P., Rossi, R. V., Canale, G. R., Melo, F. R., … Hrbek, T. (2021). An integrative analysis uncovers a new, pseudo-cryptic species of Amazonian marmoset (Primates: Callitrichidae: *Mico*) from the arc of deforestation. Scientific Reports 11(1), 1–13.

Cortés-Ortiz, L., Bermingham, E., Rico, C., Rodrıguez-Luna, E., Sampaio, I., & Ruiz-Garcıa, M. (2003). Molecular systematics and biogeography of the Neotropical monkey genus, Alouatta. Molecular Phylogenetics and Evolution 26(1), 64–81.

Cracraft, J. (1985). Historical biogeography and patterns of differentiation within the South American avifauna: areas of endemism. Ornithological Monographs 49–84.

Eberhard, J. R., & Bermingham, E. (2005). Phylogeny and comparative biogeography of Pionopsitta parrots and Pteroglossus toucans. Molecular Phylogenetics and Evolution 36(2), 288–304.

Fernandes, A. M., Wink, M., & Aleixo, A. (2012). Phylogeography of the chestnut-tailed antbird (Myrmeciza hemimelaena) clarifies the role of rivers in Amazonian biogeography. Journal of Biogeography 39(8), 1524–1535.

Figueiredo-Vázquez, C., Lourenço, A., & Velo-Antón, G. (2021). Riverine barriers to gene flow in a salamander with both aquatic and terrestrial reproduction. Evolutionary Ecology 35(3), 483–511.

Fordham, G., Shanee, S., & Peck, M. (2020). Effect of river size on Amazonian primate community structure: a biogeographic analysis using updated taxonomic assessments. American Journal of Primatology 82(7), e23136.

Gascon, C., Malcolm, J. R., Patton, J. L., da Silva, M. N., Bogart, J. P., Lougheed … Boag, P. T. (2000). Riverine barriers and the geographic distribution of Amazonian species. Proceedings of the National Academy of Sciences 97(25), 13672–13677.

Harcourt, A. H., & Wood, M. A. (2012). Rivers as barriers to primate distributions in Africa. International Journal of Primatology 33(1), 168–183.

Hayes, F. E., & Sewlal, J. (2004). The Amazon River as a dispersal barrier to passerine birds: effects of river width, habitat and taxonomy. Journal of Biogeography 31(11), 1809–1818.

Hazzi, N. A., Moreno, J. S., Ortiz-Movliav, C., & Palacio, R. D. (2018). Biogeographic regions and events of isolation and diversification of the endemic biota of the tropical Andes. Proceedings of the National Academy of Sciences 115(31), 7985–7990.

Helenbrook, W. D., & Valdez, J. W. (2021). Species distribution and conservation assessment of the black-headed night monkey (*Aotus nigriceps*): a species of Least Concern that faces widespread anthropogenic threats. Primates 62: 817–825.

Kopuchian, C., Campagna, L., Lijtmaer, D. A., Cabanne, G. S., García, N. C., … et. al. (2020). A test of the riverine barrier hypothesis in the largest subtropical river basin in the Neotropics. Molecular Ecology 29(12), 2137–2149.

Janiak, M. C., F. E. Silva, R. M. D. Beck, D. de Vries, L. F. K. Kuderna, N. S. Torosin, A. D. Melin, T. Marquès-Bonet, I. B. Goodhead, M. Messias, M. N. F. da Silva, I. Sampaio, I. P. Farias, R. Rossi, F. R. de Melo, J. Valsecchi, T. Hrbek and J. P. Boubli (2022). Two hundred and five newly assembled mitogenomes provide mixed evidence for rivers as drivers of speciation for Amazonian primates. Molecular Ecology 31(14): 3888–3902.

Lima, M. G., eSilva-Júnior, J. D. S., Černý, D., Buckner, J. C., Aleixo, A., Chang, J., … Alfaro, J. W. L. (2018). A phylogenomic perspective on the robust capuchin monkey (*Sapajus*) radiation: First evidence for extensive population admixture across South America. Molecular Phylogenetics and Evolution 124, 137–150.

Lomolino, M. V., Riddle, B. R., Whittaker, R. J., & Brown, J. H. (2010). Biogeography (Sinauer, Sunderland, MA).

Martins-Junior, A. M. G., Carneiro, J., Sampaio, I., Ferrari, S. F., & Schneider, H. (2018). Phylogenetic relationships among *Capuchin* (Cebidae, Platyrrhini) lineages: An old event of sympatry explains the current distribution of Cebus and Sapajus. Genetics and Molecular Biology 41, 699–712.

Martins-Junior, A. M., Sampaio, I., Silva, A., Boubli, J., Hrbek, T., Farias, I., Ruiz-Garcia, M., & Schneider, H. (2022). Out of the shadows: Multilocus systematics and biogeography of night monkeys suggest a Central Amazonian origin and a very recent widespread southeastward expansion in South America. Molecular Phylogenetics and Evolution 170, 107426.

Méndez-Camacho, K., Leon-Alvarado, O., & Miranda-Esquivel, D. R. (2021). Biogeographic evidence supports the Old Amazon hypothesis for the formation of the Amazon fluvial system. PeerJ 9, e12533.

Mercês, M. P., Alfaro, J. W. L., Ferreira, W. A., Harada, M. L., & Júnior, J. S. S. (2015). Morphology and mitochondrial phylogenetics reveal that the Amazon River separates two eastern squirrel monkey species: *Saimiri sciureus* and *S. collinsi*. Molecular Phylogenetics and Evolution 82, 426–435.

Moraes, L. J., Pavan, D., Barros, M. C., & Ribas, C. C. (2016). The combined influence of riverine barriers and flooding gradients on biogeographical patterns for amphibians and squamates in south-eastern Amazonia. Journal of Biogeography 43(11), 2113–2124.

Moura, M. R., & Jetz, W. (2021). Shortfalls and opportunities in terrestrial vertebrate species discovery. Nature Ecology & Evolution 5(5), 631–639.

Musher, L. J., Giakoumis, M., Albert, J., Del-Rio, G., Rego, M., Thom, G., … Cracraft, J. (2022). River network rearrangements promote speciation in lowland Amazonian birds. Science Advances 8(14), eabn1099.

Oliveira, U., Vasconcelos, M. F., & Santos, A. J. (2017). Biogeography of Amazon birds: rivers limit species composition, but not areas of endemism. Scientific Reports 7(1), 1–11.

Naka, L. N., & Brumfield, R. T. (2018). The dual role of Amazonian rivers in the generation and maintenance of avian diversity. Science Advances 4(8), eaar8575.

Nazareno, A. G., Dick, C. W., & Lohmann, L. G. (2019). A biogeographic barrier test reveals a strong genetic structure for a canopy-emergent Amazon tree species. Scientific Reports 9(1), 1–11.

Paglia, A. P., Fonseca, G. D., Rylands, A. B., Herrmann, G., Aguiar, L. M. S., Chiarello, A. G., & Patton, J. L. (2012). Lista Anotada dos Mamíferos do Brasil Occasional Papers in Conservation Biology 6.

Patton, J. L., Da Silva, M. N. F., & Malcolm, J. R. (1994). Gene genealogy and differentiation among arboreal spiny rats (Rodentia: Echimyidae) of the Amazon basin: a test of the riverine barrier hypothesis. Evolution 48(4), 1314–1323.

Pomara, L. Y., Ruokolainen, K., & Young, K. R. (2014). Avian species composition across the A mazon River: the roles of dispersal limitation and environmental heterogeneity. Journal of Biogeography 41(4), 784–796.

Prance, G. T. (1982). A review of the phytogeographic evidences for Pleistocene climate changes in the Neotropics. Annals of the Missouri Botanical Garden 594–624.

Rocha, D. G. D., & Kaefer, I. L. (2019). What has become of the refugia hypothesis to explain biological diversity in Amazonia?. Ecology and Evolution 9(7), 4302–4309.

Ribas, C. C., Aleixo, A., Nogueira, A. C., Miyaki, C. Y., & Cracraft, J. (2012). A palaeobiogeographic model for biotic diversification within Amazonia over the past three million years. Proceedings of the Royal Society B: Biological Sciences 279(1729), 681–689.

Ruokolainen, K., Moulatlet, G. M., Zuquim, G., Hoorn, C., & Tuomisto, H. (2019). Geologically recent rearrangements in central Amazonian river network and their importance for the riverine barrier hypothesis. Frontiers of Biogeography 11(3).

Ruiz-García, M., Luengas-Villamil, K., Leguizamon, N., de Thoisy, B., & Gálvez, H. (2015). Molecular phylogenetics and phylogeography of all the *Saimiri* taxa (Cebidae, Primates) inferred from mt COI and COII gene sequences. Primates 56(2), 145–161.

Ruiz-García, M., Cerón, Á., Sánchez-Castillo, S., Rueda-Zozaya, P., Pinedo-Castro, M., … et. al. (2017). Phylogeography of the mantled howler monkey (*Alouatta palliata*; Atelidae, Primates) across its geographical range by means of mitochondrial genetic analyses and new insights about the phylogeny of Alouatta. Folia Primatologica 88(5), 421–454.

Ruiz-García, M., Albino, A., Pinedo-Castro, M., Zeballos, H., Bello, A., Leguizamon, N., & Shostell, J. M. (2019). First molecular phylogenetic analysis of the *Lagothrix* taxon living in southern Peru and northern Bolivia: *Lagothrix lagothricha tschudii* (Atelidae, Primates), a new subspecies. Folia Primatologica 90(4), 215–239.

Silva, J. M. C., Novaes, F. C. & Oren, D. C. (2002). Differentiation of *Xiphocolaptes* (Dendrocolaptidae) across the river Xingu, Brazilian Amazonia: recognition of a new phylogenetic species and biogeographic implications. British Ornithology Club 185–194.

Silva, S. M., Peterson, A. T., Carneiro, L., Burlamaqui, T. C. T., Ribas, C. C., Sousa-Neves, … Aleixo, A. (2019). A dynamic continental moisture gradient drove Amazonian bird diversification. Science Advances 5(7), eaat5752.

Turchetto-Zolet, A. C., Pinheiro, F., Salgueiro, F., & Palma-Silva, C. (2013). Phylogeographical patterns shed light on evolutionary process in South America. Molecular Ecology 22(5), 1193–1213.

Wallace, A. (1852). On the monkeys of the Amazon. Proceedings of the Zoological Society of London 20, 107–110.

Wallace, A. R. (1876). The geographical distribution of animals. Hafner, New York, 145.

